# A two-decade microbial time series from a freshwater lake, introducing the limony and TYMEFLIES datasets

**DOI:** 10.1101/2022.08.04.502869

**Authors:** Robin R. Rohwer, Katherine D. McMahon

## Abstract

How microbial communities change over time is key to understanding how ecosystems will respond to global change, and to answering fundamental questions about how microbial ecology and evolution unfold. However, our understanding of microbial change is limited by a lack of long-term observations. Using archived filters from freshwater Lake Mendota (WI, USA), we created a 20-year microbial time series that begins in year 2000, a decade before Illumina DNA sequencing hit the market. We characterized 1,023 samples and controls with 16S rRNA gene amplicon sequencing in the “limony” dataset, and 471 samples and controls with shotgun metagenome sequencing in the “TYMEFLIES” dataset. In addition to the raw sequencing data, we point users to associated data products: paired environmental data collected by the North Temperate Lakes Long-Term Ecological Research program (NTL-LTER), our R package to work with curated limony data, and metagenomic assemblies and metagenome-assembled genomes (MAGs) created from the TYMEFLIES dataset. The limony and TYMEFLIES datasets, along with their paired environmental data and reference genomes, are a unique and ready-to-use resource for diverse scientists including limnologists, ecologists, evolutionary biologists, bioinformaticians, modellers, and educators.

## Background & Summary

Often considered the birthplace of limnology, Lake Mendota (WI, USA) is one of the best-studied lakes in the world. Limnological records date back to the early 1900s, and in recent decades this research has expanded to include lake microbes. Here we describe in detail the collection and sequencing of two 20-year microbial time series: a 1,023-sample 16S rRNA gene amplicon dataset we refer to as limony and a 471-sample metagenome dataset we call TYMEFLIES. We also provide background on the lake, describe an R package for working with processed limony data, and point readers to complimentary data products like environmental data and metagenome-assembled genomes (MAGs). We hope this background and practical guidance will enable other scientists to analyse limony and TYMEFLIES in creative ways that are conscious of the data’s limitations while taking advantage of its strengths.

### Lake Mendota

Lake Mendota is in southern Wisconsin, on the shores of the University of Wisconsin-Madison and the Wisconsin state capitol (Fig 1). Lake Mendota is dimictic and temperate, meaning that it has a thermally stratified water column that mixes vertically twice a year in spring and fall, and the lake freezes in the winter. In a typical year, the ice-on season lasts from late December to early April, with ice thickness reaching 40 cm (Magee et al. 2016). Vertical mixing occurs immediately after ice-off in April and as temperatures cool in October, and hypolimnetic anoxia develops during the summer stratified period (Ladwig et al. 2021). Summer surface water temperatures typically reach 25ºC, and the lake is popular for recreational activities such as fishing, sailing, and swimming. In early spring (mid-April to mid-June) a clearwater phase with typical water clarities of 7-10 m delineates a shift in phytoplankton community from diatoms to Cyanobacteria (Carey et al. 2016; Matsuzaki et al. 2020). However, Lake Mendota is eutrophic meaning that it has high concentrations of nitrogen and phosphorus, and summer water clarities of 1-2 m are punctuated by surface cyanobacterial blooms, which prevent recreation due to cyanotoxins and noxious odours (Lathrop et al. 1996; Lathrop 2007; Lathrop and Carpenter 2014; Beversdorf et al. 2015, 2017). Lake Mendota is considered a fairly typical lake, although at 39.6 km^2^ it is on the larger side, and the maximum fetch of 6 miles results in significant wave action. Lake Mendota is not unusually deep, with a maximum depth of 25 m and an average depth of 12.7 m (Brock 1985).

**Fig 1.**
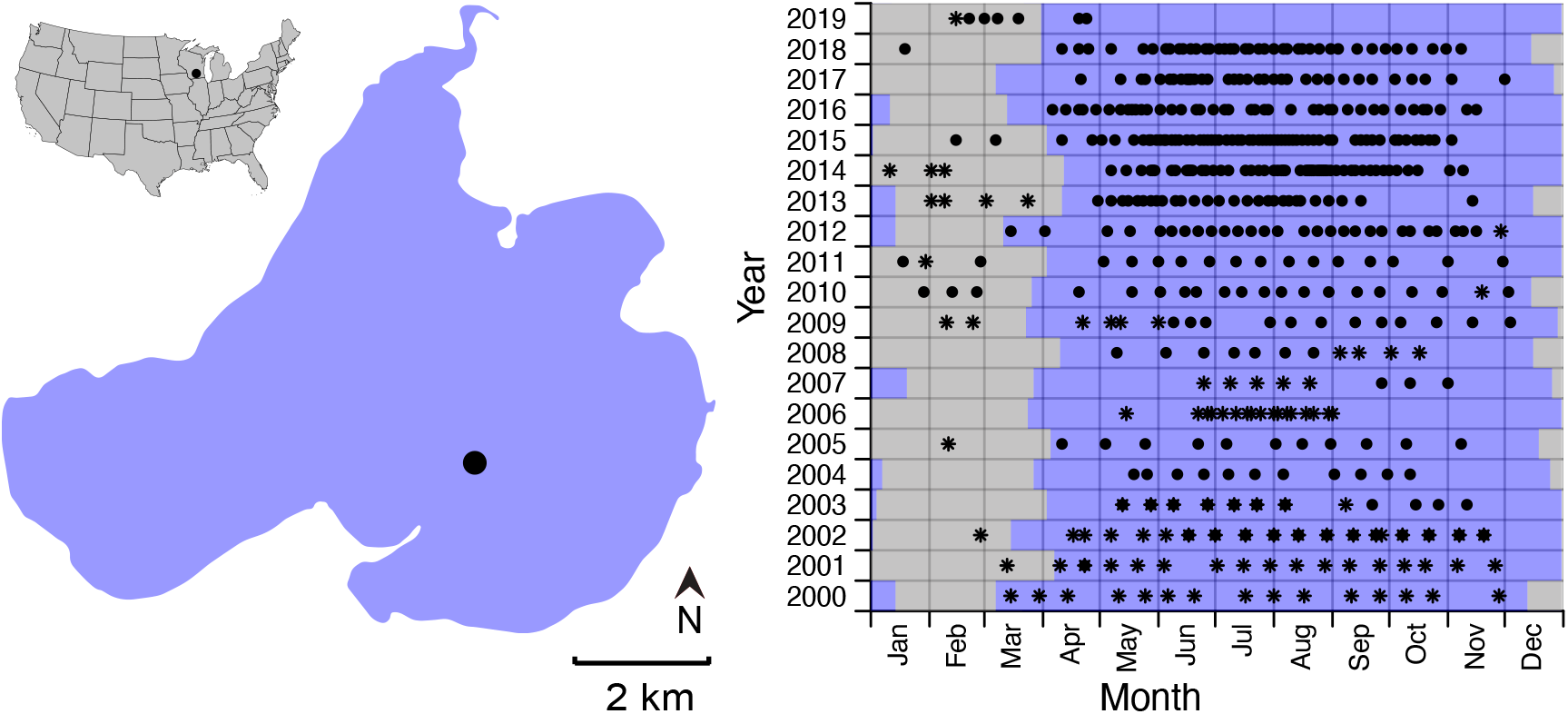
Sample dates and location. A: Map of Lake Mendota, with the deep hole sampling location indicated with a dot. B: Sample dates are displayed as points. White shading indicates periods of ice-on and blue shading indicates periods of ice-off. Samples points indicated with a star as opposed to a solid dot were collected with non-standard filtration, depth, or location (Details in Online-only Table 1). Note that some dates include both standard and non-standard samples.

Long-term research on Lake Mendota (WI, USA) began in the early 1900s (Brock 1985; Lathrop et al. 1996) and today includes research programs led by the University of Wisconsin-Madison’s Center for Limnology (CFL) and Environmental Chemistry and Technology program (EC&T), the Wisconsin Department of Natural Resources (WI DNR), the United States Geological Survey (USGS), and the North Temperate Lakes Long Term Ecological Research program (NTL-LTER). A microbial observatory (NTL-MO) (Kane 2004) was established on Lake Mendota in 1999 with funding through the “Microbial Observatories” program at the National Science Foundation, and the NTL-LTER now supports continued routine collection of microbial samples that can be used for a variety of analyses including Illumina-based 16S rRNA gene amplicon sequencing to yield “iTAGs” and shotgun sequencing to yield metagenomes. As a result of this intensive study, a wealth of long-term environmental data is publicly available through public sources such as the Environmental Data Initiative (EDI) and the USGS Water Data for the Nation.

The lake iTag measurements over nineteen years dataset, known by the acronym limony, and the Twenty Years of Metagenomes Exploring Interannual Eco/evo Shifts dataset, known by the acronym TYMEFLIES, span the duration at the time of sequencing of the archived Lake Mendota Microbial Observatory, from spring 2000 to spring 2019 (Fig 2). During this time span Lake Mendota has experienced the long-term press of climate change, observable as shorter ice cover duration, longer stratification periods, increased phosphorus loading from heavy rains, and shifting phenology (Carpenter et al. 2015; Magee et al. 2016, 2019; Matsuzaki et al. 2020; Ladwig et al. 2022). However, during the two decades covered by the limony dataset Lake Mendota has also experienced abrupt ecological shifts due to aquatic invasive species (Vander Zanden et al. 2024). In 2009, a population explosion of the invasive invertebrate predator spiny water flea (*Bythotrephes longimanus*) resulted in nearly a meter loss of water clarity due to a trophic cascade in which zooplankton predation reduced grazing pressure on phytoplankton (Walsh et al. 2016a; b, 2017, 2019), as well as shifting penology of the spring clearwater phase (Matsuzaki et al. 2020) an increase in hypolimnetic anoxia (Rohwer et al. 2024b). In the fall of 2015, invasive zebra mussels (*Dreissena polymorpha*) were first detected, and by the summer of 2016 these invasive filter feeders were already covering most hard surfaces in the near-shore zone, resulting in a 300% increase in zoobenthos and phytobenthos abundance by 2018 (Spear et al. 2022). However, zebra mussels have not resulted in clearer water despite theoretically filtering the epilimnion volume every 45 days (Spear et al. 2022), and they have resulted in increased cyanotoxins (Rohwer et al. 2023).

**Fig 2.**
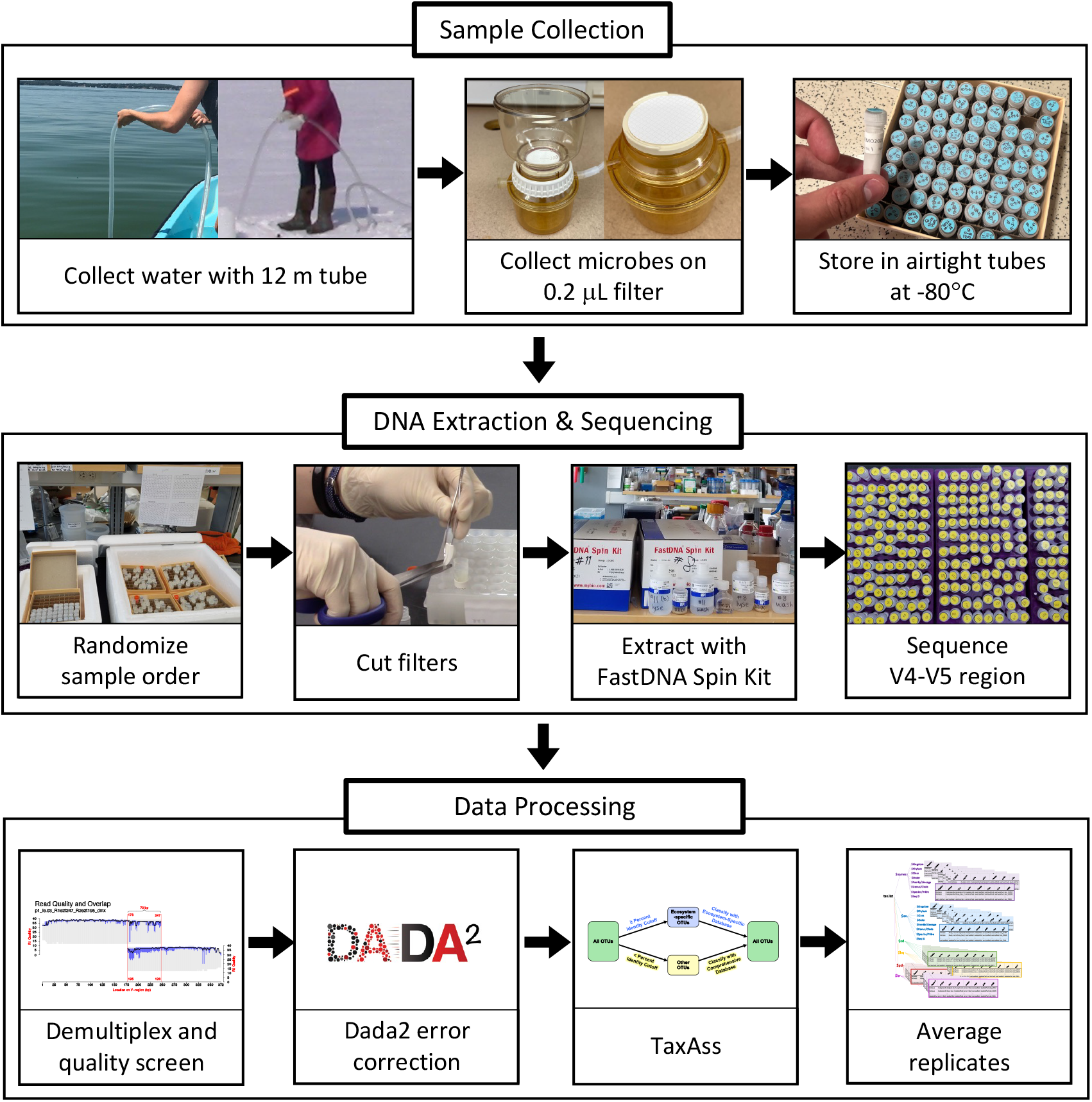
Methods overview. A schematic overview of the sampling methods, wet lab methods, and computational methods used to create the limony dataset.

The microbial community of Lake Mendota has been extensively studied prior to the limony and TYMEFLIES datasets, so broad trends about Lake Mendota’s microbial community are well established. Early studies of community composition used ARISA fingerprinting, which differentiates organisms based on the variable length of the 16S-23S rRNA gene intergenic spacer region (Fisher and Triplett 1999). These ARISA-based studies found consistent seasonal changes in bacterial community composition (Yannarell et al. 2003; Shade et al. 2007; Kara et al. 2013). Analyses of early Illumina 16S rRNA gene amplicon sequencing confirmed the strong seasonality of many Lake Mendota microbes, and identified biotic interactions as likely drivers of community change (Hall et al. 2017; Herren and McMahon 2017). In limony and TYMEFLIES, microbial phenology remains a major dynamic, with seasonal patterns observed in bacteria (Rohwer et al. 2023), including at the strain level (Rohwer et al. 2024a), as well as among protists (Krinos et al. 2024) and viruses (Zhou et al. 2024). Overlaying these short term patterns, the limony and TYMEFLIES datasets have identified long-term shifts concurrent with species invasions (Rohwer et al. 2023; Krinos et al. 2024) and environmental extremes (Rohwer et al. 2024a). Given the complexity of these datasets and the breadth of available environmental data, many more microbial trends and drivers likely yet to be discovered.

### limony: 980 iTags from 477 Sample Dates

The limony dataset is unique because of the length of its time span and sampling frequency, and because most sample dates include iTags from two biological duplicates (Fig 1). During the first decade of the time series, samples were typically collected fortnightly throughout the ice-off season, and during the second decade samples were typically collected weekly. These frequent observations are valuable for documenting community succession and revealing possible interactions over time because bacterial communities turnover on short time scales. For example, in surface seawater generation times may vary from hours to several days (Bunse and Pinhassi 2017). Regardless of generation times, multi-decadal time series are needed to tease apart true long-term change from large underlying multi-scale variability (White 2019).

The existence of biological duplicates (separate water grabs) also makes limony a powerful and unique resource. Community-derived 16S rRNA gene amplicon data are often criticized for being noisy, as evidenced by the many methods that have been created to cluster, filter, and error correct it (Eren et al. 2015; Callahan et al. 2016, p. 2; Edgar 2016; Amir et al. 2017; Schloss 2020). However, environmental data are also often highly variable at multiple scales. The NTL-MO collects biological duplicates that we sequenced in parallel, so they capture both the technical noise, which is often addressed in various data processing pipelines, and the environmental noise, which is typically ignored in microbial ecology (Prosser 2010; Lennon 2011).

### TYMEFLIES: 471 metagenomes from 463 Sample Dates

The TYMEFLIES dataset is similarly unique for its length and sample density, and also because of the depth of metagenome sequencing. TYMEFLIES samples include a single replicate from all limony sample dates with enough DNA yield to perform shotgun metagenome sequencing. Several duplicated samples exist due to a repeated “generous donor” sample included on all 5 sequencing plates, and to a few instances where sampling was nonstandard and differed between replicates. TYMEFLIES metagenome sequencing depths are 80 ± 20 million reads/sample and 23 ± 6 billion bases/sample. Deep sequencing such as this improves recovery of metagenome-assembled genomes (MAGs), and it enables genomic analyses of recovered MAGs that require sufficient depth for single nucleotide variant (SNV) calling. SNV-calling, for example, allows for strain-resolved analyses that can be used to identify evolutionary processes.

## Methods

### Sample Collection

#### Collection and transport

Water samples were collected from a central point in Lake Mendota (43°05’58.2”N 89°24’16.2”W), at the same location as a weather buoy that has collected basic physiochemical measurements during ice-off season since 2006 (Fig 1). This location is the deepest part of the lake (25 m) and is over 1.4 km from shore in all directions. Integrated 12 m samples (approximating epilimnion depth) were collected by lowering a weighted 12 m tube, plugging the top of the tube to pull out a vertical section of the water column, and then mixing this depth-integrated water sample in a 2 L bottle (Fig 2). Duplicate samples were collected from opposite sides of the boat. In the winter, samples were collected from a hole drilled by an ice augur. Some limony samples deviate from this exact method, such as winter samples that were surface water grabs from closer to shore, or samples in 2006-2007 that were depth integrated based on the measured epilimnion. These non-standard samples are indicated by stars in Fig 1 and detailed in Online-only Table 1.

#### Filtering and storage

Water was transported back to the laboratory for filtering, which could take between 15 min to an hour, depending on how quickly the field work was performed and whether the samples were transported across campus by bike or by walking. Water samples were filtered onto 0.22 μm polyethersulfone filters (Pall Corporation), and the filters were placed in airtight screw-cap tubes and stored in a -80ºC freezer until DNA extraction during 2018-2019 (Fig 2). During years 2000-2003, one hundred and nine samples were collected onto 0.22 μm filters after passing through a 10 μm prefilter. These non-standard samples are indicated by stars in Fig 1 and detailed in Online-only Table 1.

### DNA Extraction

#### Duplicates

Two filters were sequenced per sample date, one from each biological replicate (duplicate water grab). Eighty-eight sample dates deviate from this scheme however. For forty-six of these samples, two filters collected from the same water grab were used as duplicates. The remaining forty-two samples lack replicates altogether because additional filters were not available. These non-standard samples are detailed in Online-only Table 1.

#### Randomization

Filters were put in random order before performing DNA extractions and sequencing (Fig 2). Duplicates were split apart into two groups, each group’s order was randomized separately, and extractions were performed in the order of group 1 followed by group 2. For iTag analysis, each group was spread over two sequencing plates, so that duplicates were sequenced in separate Illumina runs. For metagenome analysis, the replicate with higher DNA yield was chosen and the random order was generally maintained. Samples were never ordered by date, but for example low-yield samples were grouped into the same sequencing run which was subject to manual pipetting at JGI. Extraction and sequencing orders are detailed in Online-only Table 1.

#### Extractions

DNA extractions were performed by hand using FastDNA spin kits with Lysing Matrix A used for bead-beating (MP Biomedicals) (Fig 2). To improve cell lysis during bead beating, folded filters were cut into strips using flame-sterilized scissors and tweezers, and only half of each filter was included in bead beating and DNA extraction. In preliminary testing, we found we obtained the same or higher yield from half filters, which we attributed to less-full sample tubes improving the efficacy of bead beating. The first and last extraction of every 100-extraction kit was performed on a sterile filter as a blank. All extractions were performed by the same person (RRR) according to the same method, and no differences in DNA yield were observed by extraction date or kit. Extraction yields are provided in Online-only Table 1.

### Creating the limony dataset

#### 16S rRNA gene amplicon sequencing

Sequencing was performed at Argonne National Lab according the procedure established by Walters *et al*. (2016) and recommended by the Earth Microbiome Project. The V4-V5 variable region was amplified (515F-Y GTGYCAGCMGCCGCGGTAA, 926R CCGYCAATTYMTTTRAGTTT), and sequencing was performed on an Illumina MiSeq instrument using version 2 chemistry. Sequencing was split across 4 plates that multiplexed 245, 247, 269, and 262 samples each. The sequencing plate for each sample is provided in Online-only Table 1.

#### Quality filtering and demultiplexing

Before demultiplexing, index reads were quality filtered using vsearch (Rognes et al. 2016) (vsearch v2.13.3) to retain only index reads where the expected errors were **≤** 0.03 (`--fastq_maxee = 0.03`). Mothur was then used to remove the matching sequences for the forward and reverse read files (Schloss 2020) (mothur v.1.44.1) (`list.seqs()` and `get.seqs()`). Quality control of index reads can remove cross-contamination that occurs during sequencing, sometimes called cross-talk or index hopping. This is not typically considered a major concern with the MiSeq platform (Larsson et al. 2018; van der Valk et al. 2018), but it has been observed at low levels ranging from 0.16 - 0.2% (Nelson et al. 2014; D’Amore et al. 2016). We chose a maximum expected errors cut-off of 0.03, to approximate the mean quality score cutoff of 25 suggested by Wright and Vestigian (2016).

Read quality cut-offs were chosen for each of the four limony sequencing plates separately. Cut-offs for Read quality were chosen by running vsearch (Rognes et al. 2016) with a range of expected error cutoffs and truncation lengths (`--fastq_trunclen`), and choosing a cutoff of acceptable quality and length that retained the most sequences. A maximum expected errors of 2 (`--fastq_maxee = 2`) was applied to all forward and reverse reads following truncation of the forward reads at 247 bp (plates 1 and 2) or 250 bp (plates 3 and 4) and truncation of the reverse reads at 195 bp (plates 1 and 2) or 192 bp (plates 3 and 4). Sequences were demultiplexed using qiime (Estaki et al. 2020) (qiime2-2019.4) without golay correction of the barcode sequences.

#### Error correction using dada2

For dada2 error correction (dada2 version 1.12.1) (Callahan et al. 2016), learning errors and pooled error correction were run separately on each plate, as recommended by the dada2 big data workflow tutorial. After removing phiX and merging forward and reverse reads, the four plates were combined for chimera detection.

#### Taxonomy assignment using TaxAss with the FreshTrain and Silva

We assigned taxonomy with the Silva database (Quast et al. 2013) (silva138) and the FreshTrain database (Newton et al. 2011) (FreshTrain15Jun2020silva138) using the TaxAss workflow (Rohwer et al. 2018) (v2.0.0) with a 98 percent identity cut-off. In addition to classifying freshwater heterotrophs with the FreshTrain nomenclature, TaxAss also adjusts the names in the Silva database to make duplicate names unique, and it renames the unknown sequences with the uniform format of `unnamed.NearestParentName`. Unique names within and between levels are required by our limony R scripts. Additionally, *Cyanobacteria* nomenclature from silva138 was manually edited to remove strain names from the genus-level assignments. These strain names were instead indicated at the species level, a level that is not included in the Silva database. The edited *Cyanobacteria* nomenclature is available on the TaxAss GitHub repository (https://github.com/McMahonLab/TaxAss).

#### Removing samples with low sequencing depth

The distribution of reads per sample approximated a normal distribution, and we chose a cut-off that maximized the number of samples kept, prioritizing the existence of duplicates over sequencing depth. A cut-off of 6000 reads per sample removed only 10 samples and maintained the resolution of a single read at 0.017% relative abundance. We did not rarefy the data, following the recommendation of McMurdie and Holmes (2014).

#### Processing in R

Custom R scripts were used to further process the samples by averaging replicates, calculating standard deviation, calculating several metrics of quantitation limit, and propagating these calculations to different taxonomic groupings. The exact annotated scripts are available at github.com/rrohwer/limony, and users can repeat this processing with different choices about library size or abundance cut-offs. After separating controls and applying cut-offs, the remaining replicate samples were combined. Mean abundances were calculated, along with standard deviations and information about whether an OTU was present in all replicates or at an abundance below the resolution of the smallest library size. After calculating these values at the OTU level, OTUs were grouped for each taxa level. Average abundances were summed, standard deviations were calculated by error propagation, and boolean information about replication and resolution was applied to all constituents at each taxon level. The result of this processing is stored as a list of each metric, with sub-lists for each taxon level (Fig 3). More details on the processed data format are available in the Usage Notes section.

**Fig 3.**
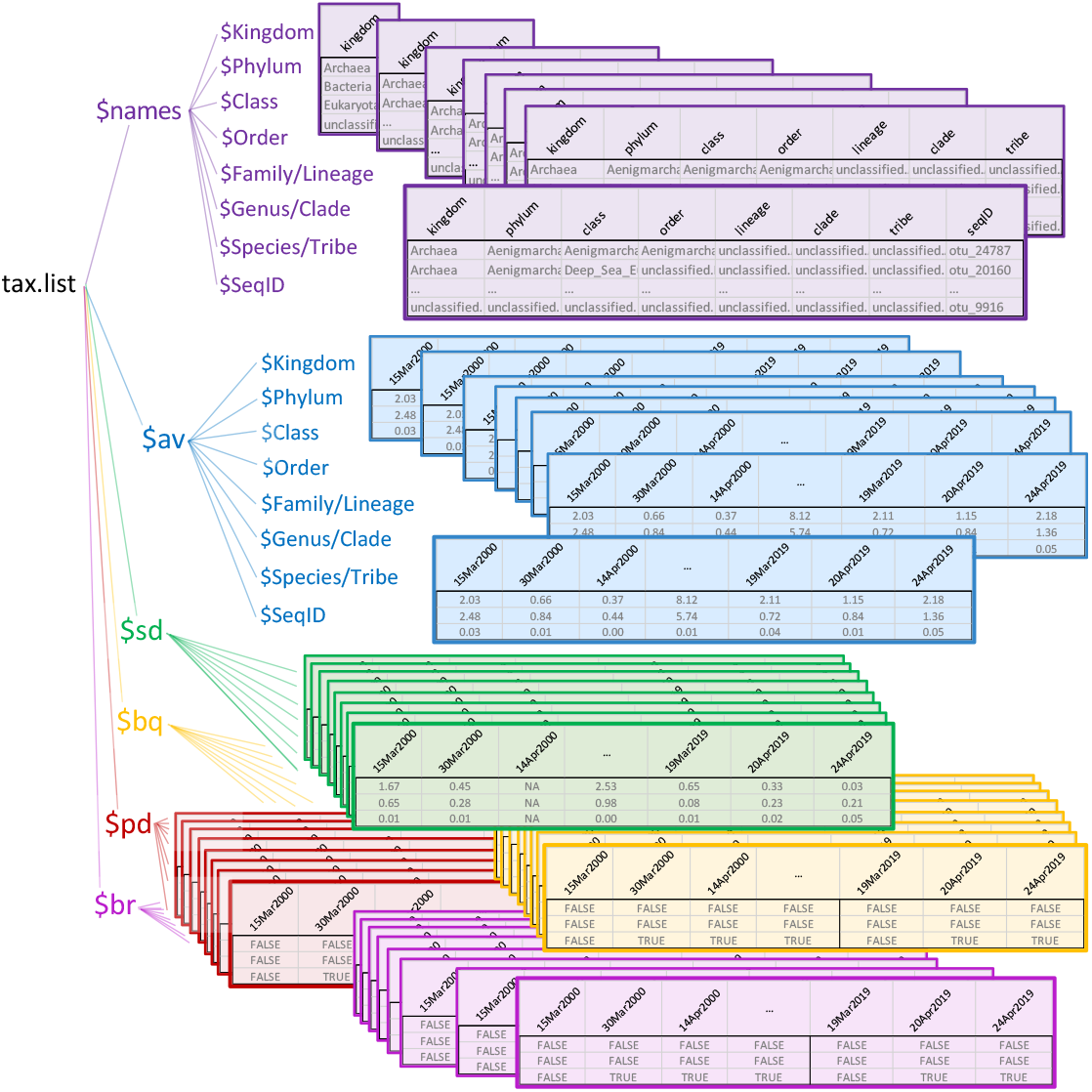
Limony nested list data structure. A visual representation of the nested list data structure in which processed data are provided as part of the limony R package. Abbreviations: av = average, sd = standard deviation, bq = below quantification, pd = partially detected, br = below resolution.

### Creating the TYMEFLIES dataset

#### Sample preparation

The DNA used for TYMEFLIES metagenome sequencing came from the same sample extractions as those used for the limony iTag dataset. However, in most cases only one biological replicate was sequenced with shotgun metagenomics. Since more DNA is required for metagenome sequencing, this replicate was typically chosen based on its higher DNA yield. Every sample date in the limony dataset with high enough DNA yield was also sent for metagenome sequencing. Online-only Table 1 can be used to link TYMEFLIES metagenomes with the matching limony iTag sample from the same DNA extraction.

#### Shotgun metagenome Sequencing

Samples were sent to the Department of Energy’s Joint Genome Institute (JGI) for shotgun metagenome sequencing. A NovaSeq 6000 Sequencing System was used with an S4 flow cell to perform paired-end shotgun sequencing. The 471 samples were sequenced to a read depth of 80 ± 20 million reads/sample (23 ± 6 billion bases/sample). Samples were randomly ordered across across 5 plates submitted with 93, 93, 93, 93, and 74 samples each, as well as 30 low-DNA tubes requiring special manual sequencing preparation. Each of these 6 submissions included a repeat generous donor control sample. The sequencing plate for each sample is provided in Online-only Table 1.

### Data Records

#### limony

Raw sequencing data for limony is available from the NCBI Sequence Read Archive under BioProject accession number PRJNA846788. A key linking sequencing codes to sample names and NCBI Run accessions (SRR#) is included as part of Online-only Table 1. The processed and curated limony dataset is available for download as an R package at github.com/rrohwer/limony.

### TYMEFLIES

Raw sequencing data for TYMEFLIES was submitted to the NCBI Sequence Read Archive by the JGI Integrated Microbial Genomes & Microbiomes (IMG/M) pipeline, so each sample has a different BioProject accession. However, the Umbrella BioProject PRJNA1056043 links all TYMEFLIES samples as well as some associated assemblies and MAGs. A key with individual BioProject numbers, as well as sequencing codes, sample names and NCBI Run accessions (SRR#) is included as part of Online-only Table 1.

### Environmental

The North Temperate Lake Long-Term Ecological Research program (NTL-LTER) collects routine limnological measurements, and their data products are available through the Environmental Data Initiative (EDI) (https://environmentaldatainitiative.org).

Some environmental data collected directly alongside the microbial samples is also available from the Environmental Data Initiative (EDI) (https://environmentaldatainitiative.org). Secchi disk water clarity measurements can be found under accession number knb-lter-ntl.416.1 and water temperature and dissolved oxygen measurements can be found under accession number knb-lter-ntl.415.2 (Rohwer and McMahon 2022a; b). Data packages from EDI are easy to access and can be downloaded directly into R, python, or matlab, or downloaded as a plain text file.

### Technical Validation

#### Randomization, Controls, Mock Communities, and Duplicates

The order of sample extractions and sequencing was randomized, with limony duplicates placed on separate sequencing plates. The first and last extraction of every 100-sample prep kit was a clean filter. These 22 extraction blanks were spread across the 4 limony MiSeq plates, and can be used to detect kit contamination, cross contamination, and sequencing cross-talk. Each of the four limony sequencing plates also included 2 mock communities and 3 repeated “generous donor” samples to control for inconsistencies in community composition as observed by Yeh *et al*. (2018). The two mock communities were an even and a staggered marine mock community provided by the Fuhrman Lab (Yeh et al. 2018). A generous donor control is a repeated sample, which is less controlled than a mock community but is more representative of the actual community composition, diversity, and evenness. Three generous donors were created from different seasons to encompass a range of lake communities. These were collected with the same methods as the limony samples, but more filters were extracted and the extracted DNA was mixed before being divided into aliquots. The generous donor samples are from May 15, 2014, August 26, 2014, and November 8, 2018. The TYMEFLIES sequencing included the November 8, 2018 generous donor sample as part of all 6 individual sample submissions, and sequencing plates also included the standard blank wells used by JGI’s internal quality control pipeline.

### Usage Notes

#### Sample Metadata

A table that lists sample names, sample dates, sequencing codes (as identified on NCBI SRA), and whether samples were taken with non-standard depth, filtration, or location is available as Online-only Table 1.

#### Additional Microbial Resources

A manually curated taxonomy of abundant freshwater heterotrophs was created by Newton *et al*. using full length clone libraries (Newton et al. 2011). This Freshwater Training set (the FreshTrain) enables fine-resolution taxonomy classifications of iTag data when used with a Taxonomy Assignment tool (TaxAss) to classify 16S rRNA gene amplicon data (Rohwer et al. 2018). We encourage users of limony to take advantage of this unique taxonomy resource so that their results are easily comparable to other freshwater datasets, and so that they can perform taxonomic analyses at ecologically relevant levels. Directions, scripts, and the most recent FreshTrain releases are available at github.com/McMahonLab/TaxAss.

Several processed datasets have been produced from TYMEFLIES and are available publicly. Quality filtered metagenome read files (fastq files) and single-sample assemblies were created by the JGI IMG/M pipeline and are available from their website (https://img.jgi.doe.gov) under ITS Proposal ID 504350. A set 2,800 of MAGs created by cross-mapping samples to single-sample assemblies (Mattock and Watson 2023) and de-replicating to 96 percent average nucleotide identity (Olm et al. 2017) was created by Rohwer *et al*. (2024a) and is available from NCBI under BioProject accession number PRJNA1158976. A co-assembly of all metagenome samples was also performed by JGI and is available under NCBI BioProject accession PRJNA1134257, as well as 1,800 accompanying MAGs that are available from IMG/M under taxon identifier 3300059473 (Oliver et al. 2024).

#### Available Environmental Data

Lake Mendota is part of the NTL-LTER site, which conducts routine sampling of water chemistry, lake physical characteristics, phytoplankton and zooplankton, macrophytes, and fish. These relevant datasets span the time period of the limony dataset and are all available through the Environmental Data Initiative. Users can find all available data by searching for datasets associated with the NTL site. In particular, users may be interested in the available long-term records of water temperature (Robertson 2016; Magnuson et al. 2020, 2021a; b; Rohwer and McMahon 2022a), ice cover (Robertson 2016; Magnuson et al. 2021c), water clarity (Robertson 2016; Magnuson et al. 2021d; Rohwer and McMahon 2022b), dissolved oxygen (Magnuson et al. 2021a; b; Rohwer and McMahon 2022a), and various nutrients and ions including carbon, nitrogen, and phosphorus (Magnuson et al. 2023a; b).

#### Using the limony R package

The processed limony data is available as part of an R package that can be installed using the command `devtools::install_github(“rrohwer/limony”)`. The R package is hosted at github.com/rrohwer/limony. In addition to the processed data, the R package includes scripts to easily subset the dataset by sample date and taxonomy, while maintaining the nested list data structure.

##### Nested list data structure

The processed limony data includes the taxonomic assignments, the average abundances, and the standard deviations of each OTU, as well as information about whether it was confidently observed. This information is saved in a list of matrices with matching dimensions named “av” (average), “sd” (standard deviation), “bq” (below quantification), “pd” (partial detection), and “br” (below resolution). The elements bq, pd, and br are boolean matrices, and bq is TRUE if either pd or br is TRUE. A visual schematic of this data structure is shown in Fig 3. Nested under each categorical element is a list corresponding to each taxonomic level. These levels are Kingdom, Phylum, Class, Order, Family/Lineage, Genus/Clade, Species/Tribe, and SeqID. Each level coarser than SeqID has been grouped, so that all categorical elements have been recalculated. The average of a taxon group is the sum of all daughter group averages, and the standard deviation is the propagated error of sums of the daughter standard deviations. The boolean elements bq, pd, and br are TRUE if all of their daughter constituents are TRUE.

##### Subsetting the nested list

The limony package includes functions to subset samples and taxa while maintaining the nested list structure. For example, users can subset particular sample dates, and they can exclude non-standard samples from analysis. Additionally, users can subset a particular taxonomic group of interest. Alternately, If users prefer to work with a simple OTU table, this can be obtained by subsetting the list. Examples of all of these workflows are available in the limony R package vignette, which can be viewed without installation on the limony GitHub repository (github.com/rrohwer/limony). In addition to the vignette examples, documentation is provided for all limony package functions.

## Supporting information

Online-only Table 1

## Code Availability

Code used to create the processed limony data is available at github.com/rrohwer/limony along with the limony R package.

## Acknowledgements

The maintenance of a long-term observatory requires continuous support from numerous personnel, including professors, postdoctoral fellows, technicians, graduate students, and undergraduate students. We thank all of these researchers, including sampling leads: Angela Kent, Tony Yannarell, Ashley Shade, Stuart Jones, Ryan Newton, Georgia Wolfe, Emily Kara Read, Todd Miller, Lucas Beversdorf, James Mutschler, and Riley Hale. We also thank the initial Microbial Observatory PI Eric W. Triplett.

Maintaining a long-term observational study is not possible without continuous funding. We thank all of these funding agencies for making this dataset possible:

- U.S. National Science Foundation Postdoctoral Research Fellowship in Biology, award number 2011002.
- E. Michael and Winona Foster WARF Wisconsin Idea Fellowship, 2018.
- National Institute of Food and Agriculture, U.S. Department of Agriculture, award number 2016-67012-24709.
- National Institute of Food and Agriculture, U.S. Department of Agriculture, Hatch Projects WIS01516, WIS01789, WIS03004.
- U.S. National Science Foundation North Temperate Lakes Long-Term Ecological Research site, NTL-LTER, award numbers DEB-9632853, DEB-0217533, DEB-0822700, and DEB-1440297.
- U.S. National Science Foundation Microbial Observatories program, award numbers MCB-9977903 and DEB-0702395.
- U.S. National Science Foundation INSPIRE award, DEB-1344254.

Finally, we thank Joshua J. Hamilton for securing funding for amplicon sequencing (U.S. Department of Agriculture award number 2016-67012-24709) and the DOE Joint Genome

Institute for metagenome sequencing through CSP 504350. We also thank the Fuhrman Lab for generously supplying their mock communities.

## Author contributions

- RRR maintained the fieldwork from 2013 – 2019, extracted the DNA, applied quality control and pipeline processing, wrote the scripts for custom data processing, created the R package, created the figures, and wrote the manuscript.
- KDM maintained the fieldwork from 2003 – 2019, advised the sample preparation and data processing, secured funding, and edited the manuscript.

## Supplemental Materials

**Online-only Table 1. Sample metadata**. This table links sample names with limony and TYMEFLIES sample IDs and NCBI SRA accessions, as well as with metadata such as sample dates and notes about any non-standard locations, depths, or filtration.

